# Transcriptional perturbation effects remain structured beyond the leading global trend

**DOI:** 10.64898/2026.10.05.756839

**Authors:** Qiyuan Liu, Qirui Zhang, Yilong Chen, Jinhong Du, Siming Zhao, Jingshu Wang

**Affiliations:** Department of Statistics, University of Chicago, Chicago, IL, USA; Department of Biomedical Data Science, Dartmouth College, Hanover, NH, USA; Institute of Data Science, The University of Hong Kong, Hong Kong SAR, China; Department of Statistics and Actuarial Science, The University of Hong Kong, Hong Kong SAR, China; Dartmouth Cancer Center, Lebanon, NH, USA

## Abstract

Prior work has described a global transcriptional response to gene perturbation, with many perturbations moving cells along a common direction. We find that perturbations contribute differently to this leading global trend and that substantial organization remains beyond it, including transferable perturbation-specific effects across datasets and a common hierarchy of responsive genes across perturbations. These patterns make simple strategies surprisingly competitive even in predicting effects beyond the leading global component.

---

Single-cell CRISPRi screens measure genome-wide transcriptional responses to targeted gene repression, providing a scalable view of gene function. Increasingly sophisticated models have also been developed to predict perturbation responses, although their gains vary across datasets and evaluation metrics^1–3^. At the same time, analyses of these screens have identified broad systematic responses in which many perturbations shift cells along a common transcriptional direction^1^. Beyond this leading global trend, however, context-dependent and perturbation-specific functional effects can coexist with broader organization across perturbations and cellular contexts, making it unclear how these different forms of structure combine in shaping perturbation responses. We analyze 12 large-scale CRISPRi screens with broad perturbational coverage, including several genome-wide screens, to characterize this structure and use prediction tasks to quantify how much information it carries.

We first examined how much perturbation-specific information transfers across datasets. Seven screens were from immortalized and five from pluripotent cell lines^4–10^ (Supplementary Table 1). To improve cross-dataset comparability, we focused on stronger perturbations and used noise-corrected estimates of underlying effect similarity and signal energy (Methods). Perturbation effects were most similar between screens of the same cell line, but substantial similarity remained across cell lines (Fig. 1a). We therefore asked whether this similarity could largely be explained by the leading global trend. We summarized this trend using the first singular direction in each dataset, referring to effects before and after its removal as total and residual effects, respectively. The leading trend accounted for a larger share of signal energy in immortalized screens. Importantly, perturbations did not contribute equally: their loading magnitudes were associated with pan-essentiality across DepMap cell lines^11^, particularly in immortalized screens (Fig. 1b). This suggests that the trend partly reflects fitness-related responses, consistent with prior links between systematic variation and cell-cycle and stress-response programs^1^. Yet residual effects retained substantial cross-dataset correspondence: similarity decreased mainly among immortalized screens, while changing little among pluripotent screens or between the two groups (Fig. 1c). Broader perturbation inclusion and concordant leading-factor directions across datasets further supported these conclusions (Fig. S1). We next asked how much predictive information this cross-dataset similarity carried. For each test perturbation, we predicted its target-dataset effect as a weighted average of measurements of that perturbation from other cell lines, with weights based on cross-dataset similarity estimated from training perturbations. We compared it with PRESAGE^12^ and a cross-dataset linear model^2^, both of which use low-dimensional perturbation embeddings learned from other cell lines, and with representative predictors trained within the target dataset^2,13,14^ (Methods). To capture complementary aspects of prediction, including perturbation-specific accuracy and recovery of structure beyond the leading shared trend, we evaluated weighted MSE^3^, the perturbation discrimination score (PDS)^10,15^, and residual cosine similarities (Methods). Despite its simplicity, weighted average was surprisingly competitive on several metrics. It performed best for PDS and was particularly strong for residual cosine among the most differentially expressed outcomes (Fig. 1d; Fig. S2). Thus, cross-cell-line similarity carries substantial predictive information beyond the leading global component. Moreover, the strong performance of direct weighted transfer suggests that transferable information in full source response profiles may not be easily compressed into low-dimensional embeddings.

**Figure 1.**
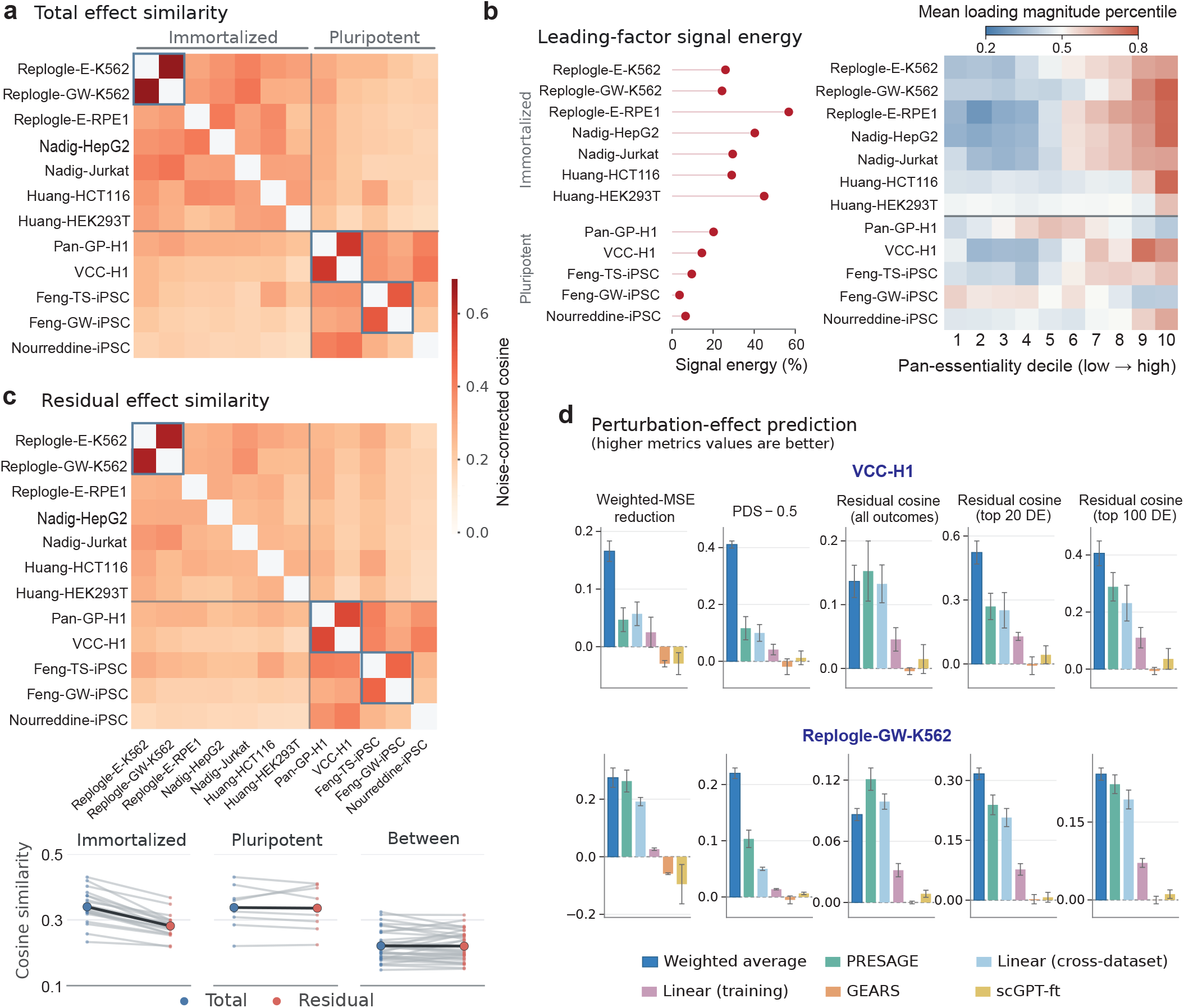
Cross-dataset structure and transferability of transcriptional perturbation effects. a.

Noise-corrected total-effect cosine similarity across datasets for shared perturbations; blue outlines indicate same-cell-line pairs. **b,** Noise-corrected leading-factor signal-energy fraction (left) and mean loading-magnitude percentile across pan-essentiality deciles (right). **c,** Noise-corrected residual-effect similarity after leading-factor removal; lower panel summarizes total and residual similarities by dataset-pair group. **d,** Perturbation-effect prediction in VCC-H1 and Replogle-GW-K562; cross-dataset methods exclude the target cell line. Zero denotes the training-mean baseline. Bars, split means; error bars, *±*1 s.d.

To further characterize perturbation responses beyond the leading global trend, we next examined how responses were organized across perturbations within each screen. We first considered patterns of statistically significant responses. Genes frequently responsive among training perturbations captured a large fraction of discoveries for held-out perturbations, revealing a shared hierarchy in which some outcome genes were consistently more responsive across perturbations. This hierarchy remained apparent after removing the leading global trend, despite a lower overall discovery density. (Fig. 2a; Fig. S3ab). Among broadly responding perturbations, a simple perturbation-agnostic differential-expression frequency (DE frequency) ranking was among the strongest predictors of both total- and residual-effect discoveries, showing that much of the ranking can be achieved without using the perturbed gene’s identity or function (Fig. 2b; Fig. S3c). Consistent with these results, an approach incorporating both DE frequency and cross-dataset weighted transfer ranked third overall in the 2025 Virtual Cell Challenge^10^.

**Figure 2.**
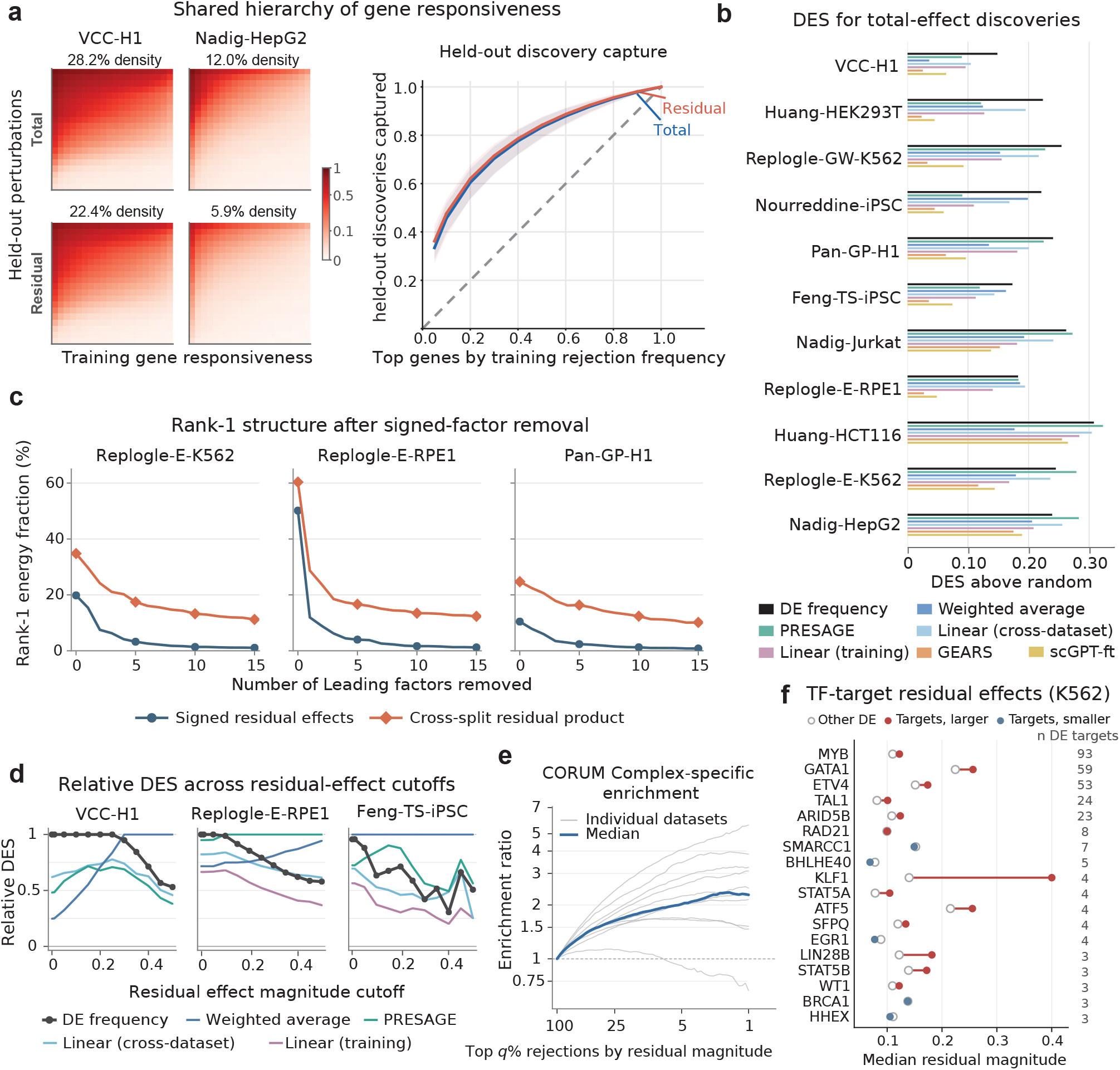
Cross-perturbation organization of residual discoveries and effects. a.

Total- and residual-effect discovery matrices (left) and held-out discovery capture across 10 datasets (right). Colors show rejection density per 20 *×* 20 bin. Heatmaps include perturbations with *≥* 20 total-effect discoveries. Blue/red curves show average total/residual capture; shading indicates 10th–90th percentiles across datasets. **b,** Differential Expression Score (DES) above the random-ranking baseline for predicting total-effect discoveries in perturbations with *≥* 20 total-effect discoveries. **c,** Rank-1 energy of signed residual effects and crosssplit residual products after removing 0–15 leading signed factors, averaged over five 50:50 data splits. **d,** Relative DES for discoveries above increasing residual-effect magnitude cutoffs, normalized to the bestperforming method at each cutoff. **e,** CORUM complex-specific enrichment among discoveries ranked by residual-effect magnitude; values *>* 1 indicate enrichment. Gray lines, individual datasets; blue, median. **f,** Median absolute residual effects for DE targets from K562-activated SCENIC+ eRegulons versus other DE genes among K562 transcription-factor perturbations with *≥* 3 DE targets. Red/blue indicate larger/smaller target medians; numbers indicate DE target counts.

The shared gene-response hierarchy suggested that residual effects might remain organized in magnitude even when signed effects differ. We therefore progressively removed leading signed factors and compared low-rank structure in signed residuals and residual magnitude. Because directly observed magnitudes are noise-biased, we used elementwise products of residual effects from independent data splits, whose expectation reflects squared true residual-effect magnitude (Methods). As additional signed factors were removed, rank-1 structure in the signed residual effects was nearly exhausted, whereas the cross-split residual-product matrices retained roughly 5-15% rank-1 energy in most datasets (Fig. 2c; Fig. S5a). Thus, substantial low-rank organization persists in residual-effect magnitude even after comparable signed structure has largely disappeared. The persistence of this magnitude structure did not imply that the shared response hierarchy was equally informative across all responses. Perturbations with few discoveries showed weaker DE-frequency performance, relatively stronger performance of direct cross-dataset transfer, and larger detected effects, suggesting that this hierarchy might be less informative for larger responses (Fig. S4). Indeed, among discoveries above progressively larger residual-effect magnitude cutoffs, DEfrequency prediction generally became less competitive than methods retaining perturbation-specific information (Fig. 2d; Figure S5b). Using CORUM protein complexes^16^, we defined complex-specific response genes as outcome genes outside the complex whose rejection frequency was selectively elevated among perturbations targeting members of that complex (Methods). These responses were generally more enriched among larger residual effects (Fig. 2e; Fig. S5c). Consistently, among DE responses to transcription factor (TF) perturbations, genes annotated by cell-type-activated SCENIC+ eRegulons^17^ as targets of the perturbed TF often showed larger residual-effect magnitudes than other DE genes from the same perturbation (Fig. 2f; Fig. S5d). These findings suggest that perturbation-specific functional organization becomes more apparent among larger responses.

Together, these results support an empirical architecture in which transcriptional perturbation responses combine broad signed variation, persistent organization of response magnitude, and perturbation-gene-specific effects. Functional organization becomes more apparent among larger residual effects, while substantial broader organization remains. These broad structures help explain why simple prediction methods can perform remarkably well, but also make it challenging to isolate and predict perturbation-specific functional responses.

## Online Methods

### Datasets and preprocessing

All datasets were processed using a common workflow based on crispyx (version 0.1.4). Cells with fewer than 100 detected genes were removed. A perturbation was retained if at least 20 remaining cells carried it; control cells were always retained. Genes detected in fewer than 100 retained cells were removed. Counts were normalized to 10^4^ per cell and log-transformed as log(1 + *x*). On the normalized log-expression scale, perturbations were excluded if the targeted gene had mean expression below 0.08 in control cells or if its expression was reduced by less than 10% in perturbed cells relative to controls. Outcome genes were excluded if their mean expression was below 0.08 in both control and perturbed cells. All analyses used perturbations and outcome genes passing these filters. The numbers of cells, perturbations, and outcome genes retained after preprocessing are reported in Supplementary Table 2.

### Perturbation-effect estimation

For each perturbation *p* and outcome gene *g* in dataset *d*, we estimated a stratum-adjusted perturbation effect by combining within-stratum differences in mean lognormalized expression. We defined strata by batch, except for the Feng datasets, where we treated each cell line-batch combination as a separate stratum.

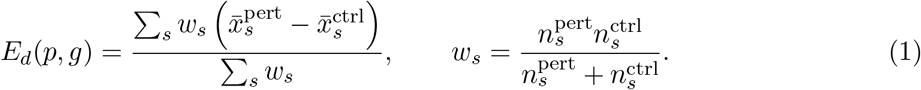

Here, 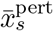 and 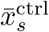 are the mean perturbed and control expressions, and 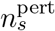 and 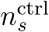 are the corresponding cell counts in strata *s*; sums are over strata containing both perturbed and control cells. This weighting gives the same estimate as a linear regression with stratum-specific intercepts and a single perturbation coefficient across strata. Collecting these estimates gives the perturbation-by-gene effect matrix *E_d_*. Standard errors SE*_d_*(*p, g*) were estimated assuming independence of cells within strata and common condition-specific variances across strata (Supplementary Methods).

### Stratified Wilcoxon tests

For each perturbation, perturbed and control cells were compared using a Wilcoxon test based on the same strata as in effect estimation. Within each stratum, we computed a Wilcoxon ranksum statistic comparing perturbed and control cells. The centered stratum-specific *U* statistics were summed across strata and standardized using their combined null variance to obtain a two-sided *P* value. The resulting *P* values were adjusted by the Benjamini-Hochberg (BH) procedure across all outcome genes retained after preprocessing, excluding the directly perturbed gene, within each perturbation.

### Strong-perturbation selection

We defined a set of strong perturbations using the following L1 perturbation-strength score. Let *q_dpg_* denote the resulting adjusted *P* -value from the BHcorrected stratified Wilcoxon tests. We defined the L1 perturbation-strength score as

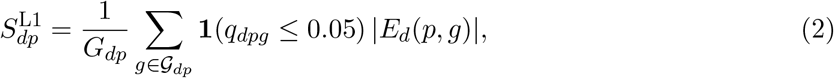

where *G_dp_* is the set of outcome genes retained after preprocessing, excluding the directly perturbed gene and *G_dp_* = *|G_dp_|*. Perturbations were ranked independently within each dataset, and we retained up to 2,000 perturbations with finite positive scores, except for VCC-H1, where up to 150 were retained. When fewer perturbations qualified, all were retained. We refer to these as strong perturbations.

### Leading global trend and residualization

For each dataset *d*, we performed an uncentered singular value decomposition (SVD) of the total-effect matrix *E_d_* using all perturbations and outcome genes, after setting each perturbation’s effect on its own target gene to zero. We represented the leading global trend by the leading singular component, 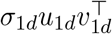, and defined the residual-effect matrix as

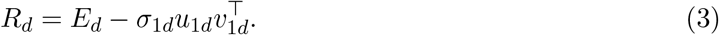

We refer to *E_d_* and *R_d_* as total and residual effects, respectively. The leading component was fitted once per dataset; subsequent perturbation and outcome-gene subsets were applied only after residualization.

### Noise-corrected cosine similarity

For each dataset pair, we compared perturbation-effect profiles for shared strong perturbations in both datasets, using outcome genes measured in both datasets and excluding the directly perturbed gene. Because estimation noise inflates squared effect norms, we corrected each norm by subtracting the summed estimated error variances:

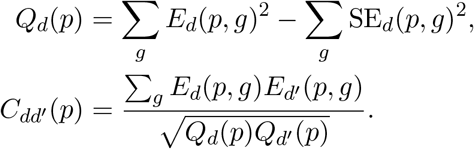

The sums are over outcome genes measured in both datasets. The correction removes the expected noise contribution to the squared norms, while the cross-dataset inner product requires no analogous correction (Supplementary Methods).

Residual-effect similarity was calculated identically after replacing *E_d_* with *R_d_*, while retaining the original standard errors. To avoid unstable cosine estimates for profiles dominated by estimation noise, a perturbation was retained only if its noise-corrected squared norm exceeded 5% of its observed squared norm in both datasets and for both total and residual effects, ensuring identical perturbation membership in the two comparisons. Retained cosine estimates were clipped to [*−*1, 1] and averaged across perturbations to obtain the dataset-pair similarity.

### Noise-corrected signal energy

Among strong perturbations in each dataset, we quantified the strength of the leading rank-one structure in the perturbation-effect signal after correcting for estimation noise. With *E_d_* restricted to these perturbations, we defined

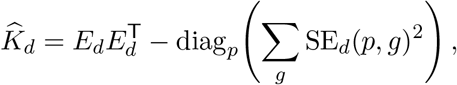

where the diagonal correction removes the estimated contribution of sampling noise to each perturbation’s squared norm. We summarized the leading rank-one signal fraction as

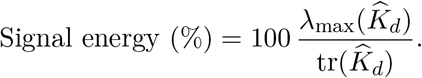

The noise correction and its justification are described further in the Supplementary Methods.

### Association with essentiality

Gene essentiality was quantified using DepMap Public 26Q1. Within each cell line, genes were ranked by Chronos gene-effect score and scaled to [0, 1], with 1 indicating the strongest dependency. For each gene, we defined panessentiality as the tenth percentile of these ranks across cell lines, so that higher values indicate consistently strong dependency across at least 90% of cell lines. The absolute leading-component perturbation loadings *|u*_1*d*_(*p*)*|* from the SVD fit to all perturbations were converted to within-dataset percentile ranks. To summarize their association with essentiality, perturbations were grouped into ten equalsized bins by the panessentiality of their target genes, and the mean loading percentile was calculated within each bin.

### Weighted-average transfer

For each target dataset, perturbations were randomly assigned to five disjoint folds, and the same folds were used for all prediction methods. Each perturbation’s target gene was excluded from all prediction metrics and discovery sets. For a target dataset, we predicted each test perturbation by a weighted average of measurements of the same perturbation in other datasets,

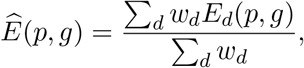

where *w_d_* was the mean cosine similarity between dataset *d* and the target across perturbations shared with the target training set, evaluated on genes measured in both datasets. For each perturbation-gene pair, only datasets from different cell types in which both the perturbation and gene were measured contributed to the average; missing measurements were ignored rather than treated as zero. Predictions were evaluated only for perturbation-gene pairs observed in at least one such dataset, and the same perturbation-gene evaluation set was used for all prediction methods. Negative weights were set to zero.

### Comparison methods

We compared the weighted average with GEARS^13^, scGPT^14^ fine-tuned on each target dataset (scGPT-ft), two versions of linearModel^2^, PRESAGE^12^, and a mean baseline that predicted each gene’s average effect across target-dataset training perturbations. Linear (training) used perturbation representations learned from the target training data, whereas Linear (cross-dataset) used representations learned from perturbation effects in other datasets. For PRESAGE, we used PRESAGE (+Perturb-seq), which augments its knowledge-source embeddings with embeddings learned from Perturbseq data. For Linear (cross-dataset) and PRESAGE, cross-dataset information was restricted to the same reference datasets available to weighted average. Further implementation details are provided in the Supplementary Methods.

### Prediction metrics

We evaluated predictions using weighted mean squared error (wMSE), PDS, and residual cosine similarity. wMSE emphasizes prediction accuracy for genes with stronger differential-expression signals by assigning them greater weight. We reported the relative reduction versus the mean baseline, (wMSE_mean_ *−* wMSE_method_)*/*wMSE_mean_, where wMSE_mean_ denotes the mean-baseline wMSE. PDS measures whether each predicted perturbation is closest in cosine distance to its corresponding observed perturbation among the test perturbations, with 1 indicating perfect discrimination (Supplementary Methods). We report PDS *−* 0.5, as the mean baseline gives the same prediction for every test perturbation and its average PDS is 0.5 and therefore corresponds to zero on this scale. For residual cosine similarity, the leading rank-one component was removed separately from the observed and predicted test-set effect matrices, and cosine similarity between corresponding residual-effect vectors was calculated over all outcome genes and, separately, over the top 20 and top 100 differentially expressed genes ranked by Wilcoxon *P* value. The mean baseline has rank-one predictions and therefore zero residual cosine similarity. Metrics were averaged over test perturbations within each fold and summarized as mean *±* s.d. across the five folds.

### Total and residual discoveries

Let *D_p_* denote the set of genes discovered for perturbation *p* by the stratified Wilcoxon analysis, using a BH-adjusted *P* value threshold of 0.05. To define residual discoveries, we required evidence both from the original Wilcoxon test and from an approximate two-sided test of the residual effect,

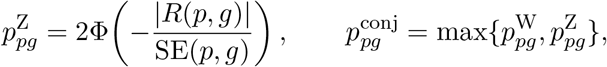

where 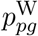 is the stratified Wilcoxon *P* value and the original standard errors were retained after residualization. The residual discovery set 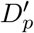 contains genes with BHadjusted 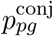 ≤ 0.05. This conjunction criterion avoids calling a gene residual-responsive based on the original differential-expression signal alone when its residual effect is weak.

### Training-set DE frequency

For discovery prediction, outcome genes were ranked within each training set by their discovery frequency across training perturbations, excluding the perturbation targeting the gene itself:

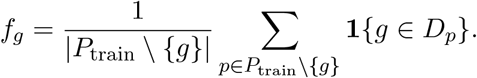

The same ranking was used for all held-out perturbations. For residual discoveries, frequencies were recomputed using 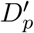 in place of *D_p_*.

### Held-out discovery structure

To assess whether the gene-responsiveness hierarchy generalized to unseen perturbations, we generated 100 random 70%*/*30% training–held-out splits within each dataset. Splits were stratified by total-effect discovery fraction to give training and held-out sets similar response-strength distributions, and the same splits were used for total and residual analyses. Within each split, genes were ranked by the corresponding training-set *f_g_*.

Held-out discoveries were summarized in two ways. Discovery heatmaps included perturbations with at least 20 total-effect discoveries, with genes ordered by the training-derived ranking. For the capture curves, we calculated the fraction of held-out discoveries captured among increasingly large fractions of the ranked genes. These analyses were restricted to the 10 datasets with sufficient discoveries for held-out evaluation. Within each dataset, capture was averaged over held-out perturbations and repeated splits, and the resulting curves were averaged equally across datasets.

### Discovery prediction

For a test perturbation *p* with *k_p_* = *|D_p_| >* 0, we took the top *k_p_* genes from each method’s ranking as the predicted discovery set 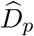 and evaluated recovery by

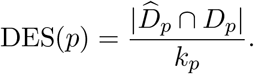

Thus, DES evaluates responsive-gene ranking at a fixed set size rather than predicting the number of differentially expressed genes. DE frequency ranked genes by *f_g_*, whereas effect-based methods ranked genes by 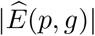; the random-ranking baseline sampled *k_p_* genes uniformly without replacement. We report DES above random as the difference between each method’s mean DES and the random-ranking baseline mean over the same perturbations. Residual DES was defined analogously using 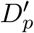 , with effect-based methods ranked by the absolute residual predicted effect after removal of the leading rank-one component.

### Signed-factor removal and residual-magnitude structure

For each dataset *d*, we progressively removed the leading signed singular components from the effect matrix *E_d_*. Suppressing the dataset index for notational simplicity, we defined 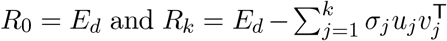. We quantified leading rank-one structure by the fraction of squared Frobenius norm captured by the leading component, 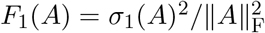 . To assess whether residual-effect magnitude contained genuine low-rank structure rather than structure induced by estimation noise, we independently estimated residual effects from two random halves of the cells and formed their elementwise product,

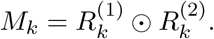

Under independent meanzero estimation errors, each entry of *M_k_* has expectation equal to the squared underlying residual effect, avoiding the additive noise-variance contribution present in squared or absolute observed effects (Supplementary Methods). For each *k*, we averaged the corresponding signed-residual and cross-split product matrices over five random cell splits before computing *F*_1_. We evaluated *k* = 0*, … ,* 15. To prevent a small number of extreme entries from dominating the leading rank-one component, entry magnitudes were capped at the 99.99th percentile before calculating *F*_1_ (Supplementary Methods).

### Discovery prediction across residual-effect magnitudes

To assess prediction performance among increasingly large residual effects, we restricted the residual discovery set for perturbation *p* to

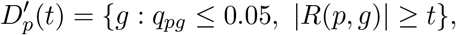

where *q_pg_* is the BH-adjusted conjunction *P* value defining residual discoveries. We evaluated *t* = 0, 0.01, and 0.05 to 0.50 in increments of 0.05, using 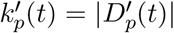 as the predicted-set size and excluding perturbations with 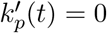. At each threshold, DE-frequency rankings were recomputed from the corresponding thresholded discovery sets among training perturbations. Effect-based methods retained their predicted residual-effect rankings and selected the top 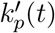 genes by absolute predicted residual effect, while the random baseline sampled 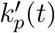 genes uniformly without replacement. Thus, the threshold was applied to the observed residual discoveries used for evaluation, rather than to the predicted effects. For each dataset and threshold, DES above random was normalized by the largest DES-above-random value among the compared methods.

### Protein-complex enrichment

Using human protein complexes from CORUM 5.0^16^, we identified outcome genes whose residual-discovery frequency was selectively elevated among perturbations targeting members of the same complex relative to perturbations targeting other complexes. Complex-specific outcome genes were required to show substantial enrichment after accounting for both perturbation-level discovery density and gene-level responsiveness, and complexes were required to contain at least three measured perturbation targets (Supplementary Methods).

We pooled residual discoveries from perturbations targeting eligible complexes, excluding perturbation– gene pairs in which the perturbed gene and outcome gene belonged to the same complex. A discovery was labeled complex-specific if the perturbed gene belonged to an eligible complex and the outcome gene was identified as complex-specific for that complex. Discoveries were ranked by residual-effect magnitude *|R*(*p, g*)*|*. At each cutoff, enrichment was defined as the fraction of complex-specific discoveries among the largest residual effects divided by their fraction among all discoveries. Enrichment curves were summarized across datasets by the median.

### Transcription-factor analyses

We used the SCENIC+ gene regulatory network inferred jointly from eight ENCODE cell lines^17^. Among these eight cell lines, K562, HepG2, and HCT116 overlapped with the cell lines represented in our perturbation datasets, and we therefore focused on these three. For each cell line, we used the cell-line-activated eRegulons reported by SCENIC+ and defined the target genes of each TF as the union of genes across its active eRegulons.

In the three datasets, Replogle-GW-K562, Nadig-HepG2 and Huang-HCT116, we analyzed perturbations of TFs represented in the corresponding cell-line-activated network. We retained TF perturbations with at least three residual discoveries among their SCENIC+ target genes and at least one residual discovery among other outcome genes. For each perturbation, we compared the median residual-effect magnitude *|R*(*p, g*)*|* among discovered TF targets with that among discovered non-target genes.

## Data availability

The single-cell perturbation datasets analyzed in this study are publicly available from the following repositories. Replogle-GW-K562, Replogle-E-K562, and Replogle-E-RPE1 are available from Zenodo (https://zenodo.org/records/10044268). Nadig-HepG2 and Nadig-Jurkat are available from GEO under accession number GSE264667. Huang-HCT116 and Huang-HEK293T are available from Figshare+ (https://doi.org/10.25452/figshare.plus.29190726.v1). VCC-H1 is available from the Arc Virtual Cell Atlas (https://github.com/ArcInstitute/arc-virtual-cell-atlas/tree/main/virtual-cell-challenge). Feng-GW-iPSC and Feng-TS-iPSC are available from Figshare (https://doi.org/10.6084/m9.figshare.27989294.v2). Nourreddine-iPSC is available from Figshare+ (https://doi.org/10.25452/figshare.plus.27261219.v1). Pan-GP-H1 is available from GEO under accession number GSE295214.

DepMap Public 26Q1 Chronos gene-effect data used to derive pan-essentiality scores were obtained from the DepMap Data Portal at https://depmap.org/portal/data_page/. CORUM 5.0 protein-complex annotations were obtained from https://mips.helmholtz-munich.de/corum/. SCENIC+ eRegulon annotations for K562, HepG2, and HCT116 used in the transcription-factor analyses were obtained from SCope at https://scope.aertslab.org/#/scenic-v2.

The preprocessed data and test statistics in the paper can be accessed through HKU DataHub (https://doi.org/10.25442/hku.34002315).

## Code availability

Code used to reproduce the analyses and figures in this study is publicly available at https://github.com/liuqiyuanhhh/perturbation-structure. Dataset preprocessing was based on crispyx (version 0.1.4), available at https://pypi.org/project/crispyx/.

## Acknowledgements

J.W. was supported in part by the National Science Foundation under grant DMS-2238656 and the National Institute of General Medical Sciences under grant R35GM162500. S.Z. and Q.Z. were supported in part by Munck-Pfefferkorn Grant Awards from the Geisel School of Medicine at Dartmouth College. J.H.D. acknowledges support from the National Natural Science Foundation of China (No. 12601487) and the Institute of Data Science at the University of Hong Kong through the HKU100 award.

Computational support was provided by the University of Chicago Research Computing Center; Dartmouth Research Computing and Data, including the Discovery Cluster; and the research computing facilities of Information Technology Services at the University of Hong Kong.

We thank Weiyi Wu and Gui Jiang of the Department of Biomedical Data Science at Dartmouth College for their support during our participation in the 2025 Virtual Cell Challenge.

## Author contributions

J.W. and S.Z. conceived the study. J.W., S.Z., and J.H.D. supervised the study. J.W. led the development of the methodology and analytical direction. Q.L. carried out the majority of the computational work, including implementation of the statistical methods, prediction benchmarking, and downstream analyses. J.H.D. developed crispyx, processed the perturbation datasets, and contributed to selected benchmarking analyses. Q.Z. performed the PRESAGE benchmarking and contributed to exploratory analyses. Y.C. contributed to the implementation and evaluation of candidate prediction methods. J.W. and Q.L. drafted the manuscript. All authors contributed to the interpretation of the results and reviewed and edited the manuscript.

