## supplements for "Transcriptional perturbation effects remain structured beyond the leading global trend"

### Supplementary Tables

| Dataset | Year | Perturbation set | Perturbations | Median cells per perturbation | Median library size |
|---|---|---|---|---|---|
| Replogle-GW-K562[1] | 2022 | Genome-wide | 9,866 | 178 | 11,304 |
| Replogle-E-K562[1] | 2022 | Essential genes | 2,057 | 121 | 13,207 |
| Nadig-Jurkat[2] | 2025 | Essential genes | 2,393 | 83 | 9,997 |
| Nadig-HepG2[2] | 2025 | Essential genes | 2,393 | 45 | 15,487 |
| Replogle-E-RPE1[1] | 2022 | Essential genes | 2,393 | 72 | 11,940 |
| Huang-HCT116[3] | 2025 | Genome-wide | 18,293 | 150 | 17,292 |
| Huang-HEK293T[3] | 2025 | Genome-wide | 18,311 | 200 | 20,699 |
| Feng-GW-iPSC[4] | 2026 | Genome-wide | 7,218 | 25 | 23,388 |
| Feng-TS-iPSC[4] | 2026 | Selected: variable effects | 444 | 1,355 | 18,614 |
| Pan-GP-H1[5] | 2026 | Selected: growth/pluripotency | 2,978 | 283 | 14,104 |
| VCC-H1[6] | 2025 | Selected: effect strength | 300 | 1,071 | 54,061 |
| Nourreddine-iPSC[7] | 2026 | Genome-wide | 11,687 | 218 | 5,119 |

Supplementary Table 1: Summary statistics of the 12 datasets before quality control. The upper block contains the seven non-pluripotent screens and the lower block contains the five pluripotent screens.

| Dataset | Perturbations | Outcome genes | Median cells per perturbation |
|---|---|---|---|
| Replogle-GW-K562 | 6,807 | 7,446 | 177 |
| Replogle-E-K562 | 1,778 | 7,646 | 122 |
| Nadig-Jurkat | 1,956 | 8,143 | 85 |
| Nadig-HepG2 | 1,813 | 8,120 | 49 |
| Replogle-E-RPE1 | 1,812 | 7,887 | 74 |
| Huang-HCT116 | 8,068 | 9,191 | 142 |
| Huang-HEK293T | 9,060 | 10,445 | 188 |
| Feng-GW-iPSC | 4,075 | 9,257 | 32 |
| Feng-TS-iPSC | 403 | 8,987 | 1,353 |
| Pan-GP-H1 | 2,164 | 8,356 | 289 |
| VCC-H1 | 293 | 9,786 | 1,077 |
| Nourreddine-iPSC | 6,507 | 9,074 | 216 |

Supplementary Table 2: Summary statistics of the 12 datasets after data preprocessing.

### Supplementary Figures

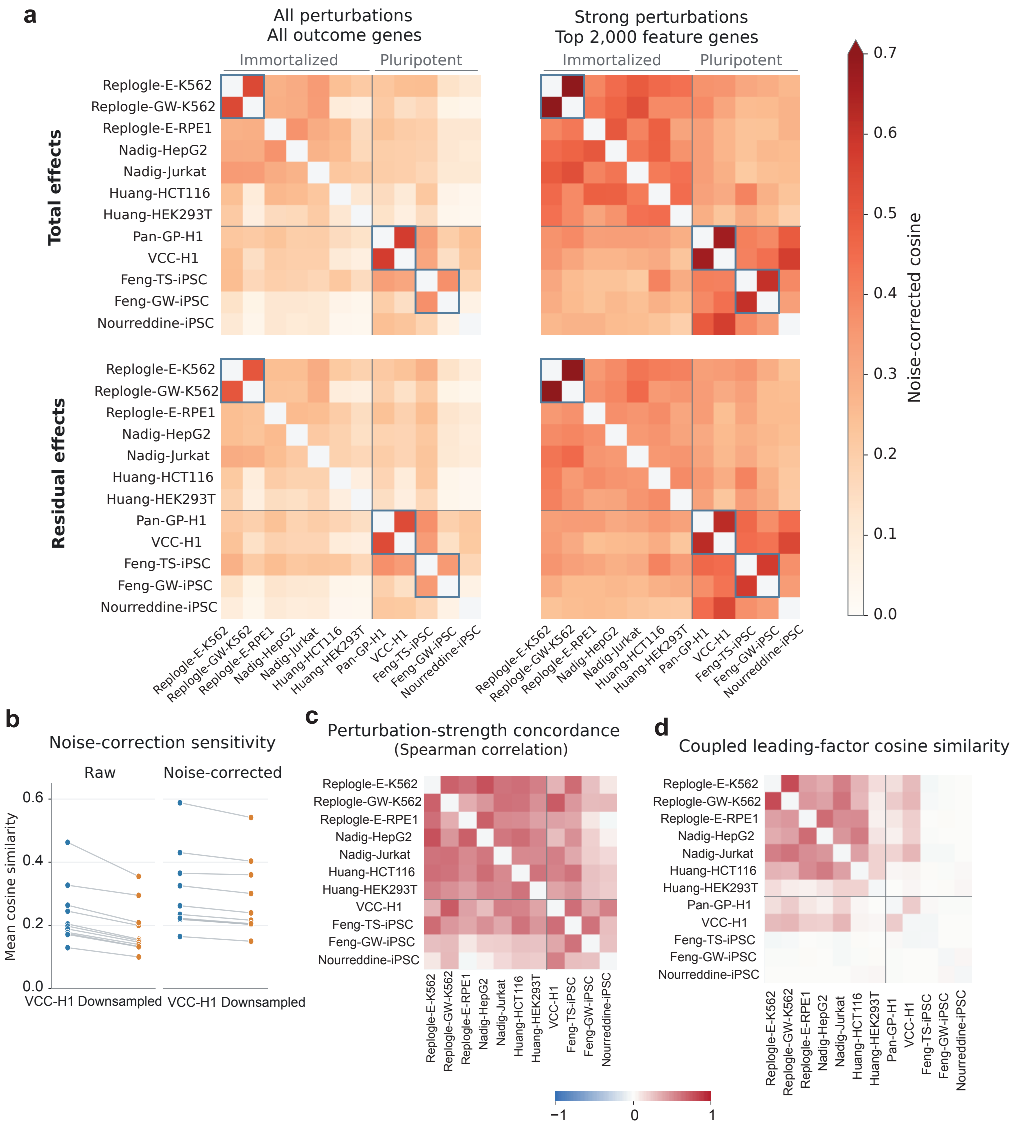


Figure S1: **Sensitivity of cross-dataset similarity and concordance of perturbation strength and leading factors across datasets. a,** Noise-corrected total-effect (top) and residual-effect (bottom) cosine similarity across datasets for shared perturbations, using all perturbations retained in preprocessing and all outcome genes (left) or strong perturbations and the top 2,000 feature genes (right). Blue outlines indicate same-cell-line pairs. **b,** Mean raw (left) and noise-corrected (right) total-effect cosine similarity between VCC-H1 and each of the other 11 datasets before (blue) and after (orange) downsampling VCC-H1 in UMI depth and cell number, over the same shared strong perturbations. Gray lines connect the same dataset. **c,** Spearman correlation of L1 perturbation-strength scores between datasets over all shared perturbations. Pan-GP-H1 is not shown because the dataset used a different perturbation selection scheme. **d,** Leading-factor concordance: cosine similarity between the leading global trend loadings of each dataset pair over shared perturbations and outcome genes.

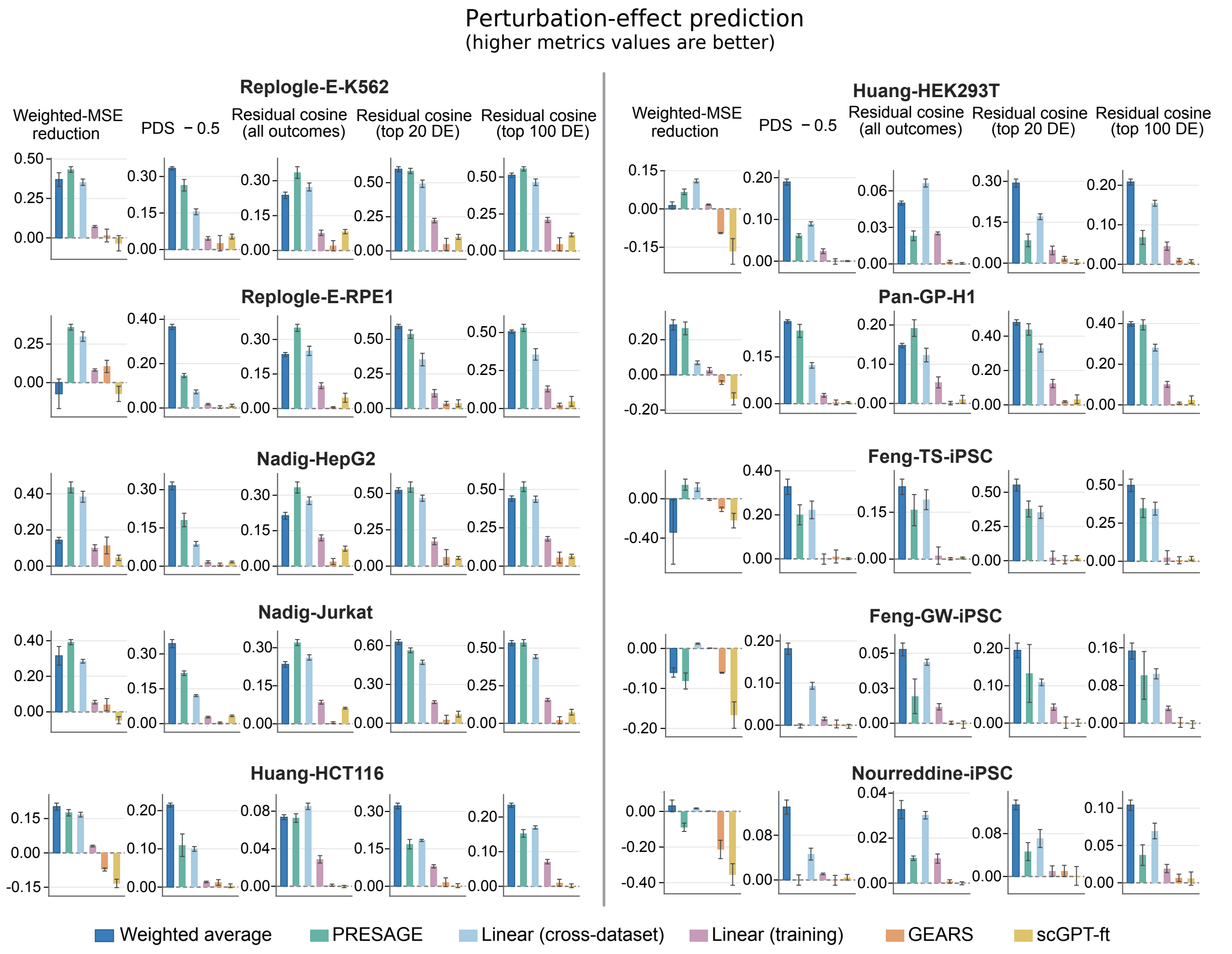


Figure S2: **Perturbation-effect prediction in the remaining 10 datasets.** Analysis as in Fig. 1d. Metrics are weighted-MSE reduction, PDS − 0.5, and residual cosine similarity over all outcome genes and over the top 20 and top 100 DE genes. Zero denotes the training-mean baseline. Bars, split means; error bars, ±1 s.d.

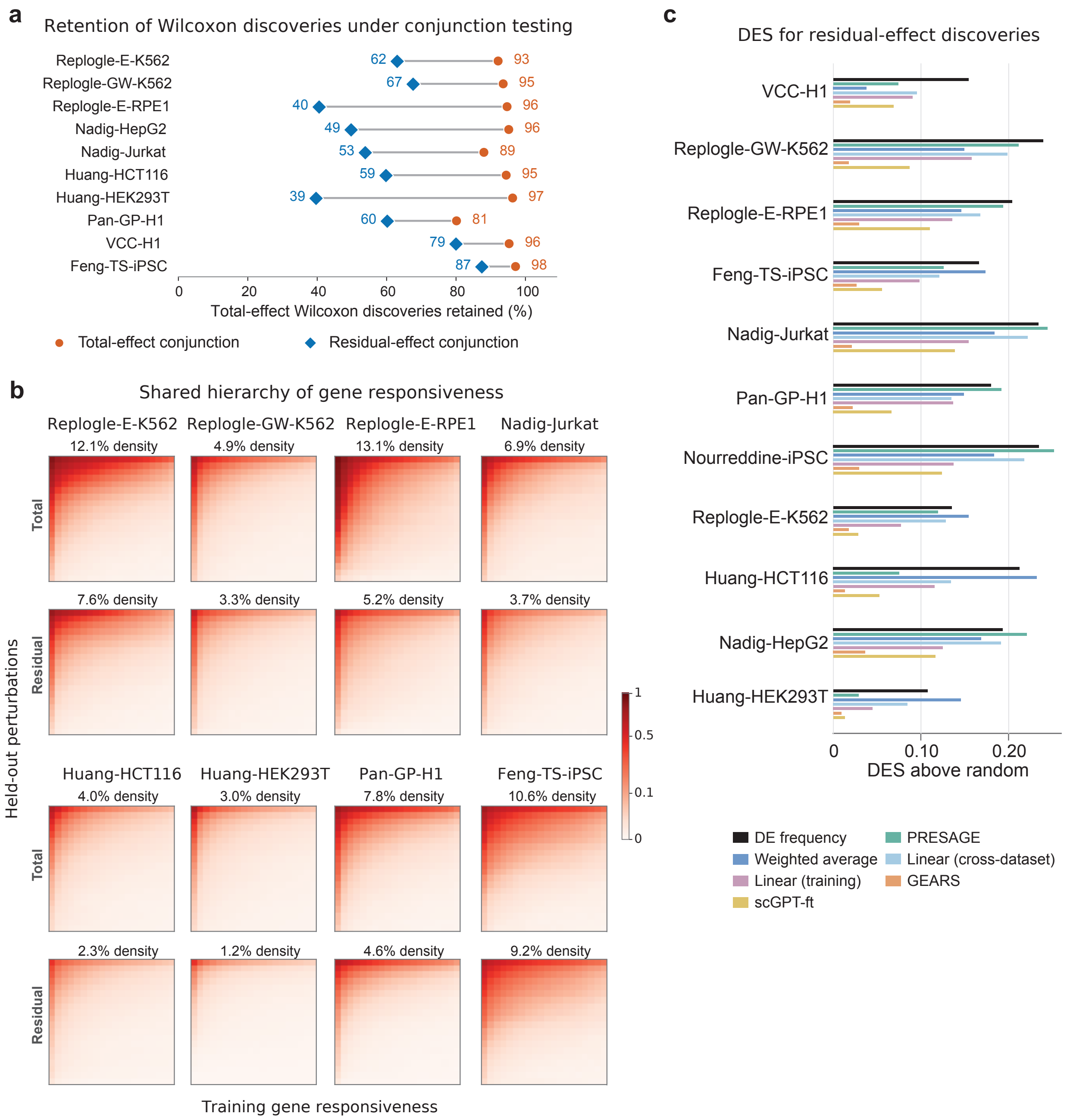


Figure S3: **Retention, cross-perturbation organization and prediction of residual-effect discoveries. a,** Percentage of total-effect discoveries retained by the conjunction criterion defining residual-effect discoveries (residual-effect conjunction) and by the same criterion applied to total effects (total-effect conjunction), in the 10 datasets in Fig. 2a, excluding the low-powered screens Feng-GW-iPSC and Norredine-iPSC. Counts are pooled over all perturbations in each dataset. **b,** Total- and residual-effect discovery matrices, as in Fig. 2a, for eight additional datasets. Held-out perturbations with $\geq 20$ total-effect discoveries (rows) are ordered by discovery fraction and outcome genes (columns) by training-set DE frequency. Colors show rejection density per $20 \times 20$ bin, averaged over 100 random 70%/30% training–held-out splits; percentages give overall discovery density. **c,** DES above the random-ranking baseline for predicting residual-effect discoveries in perturbations with $\geq 20$ total-effect discoveries.

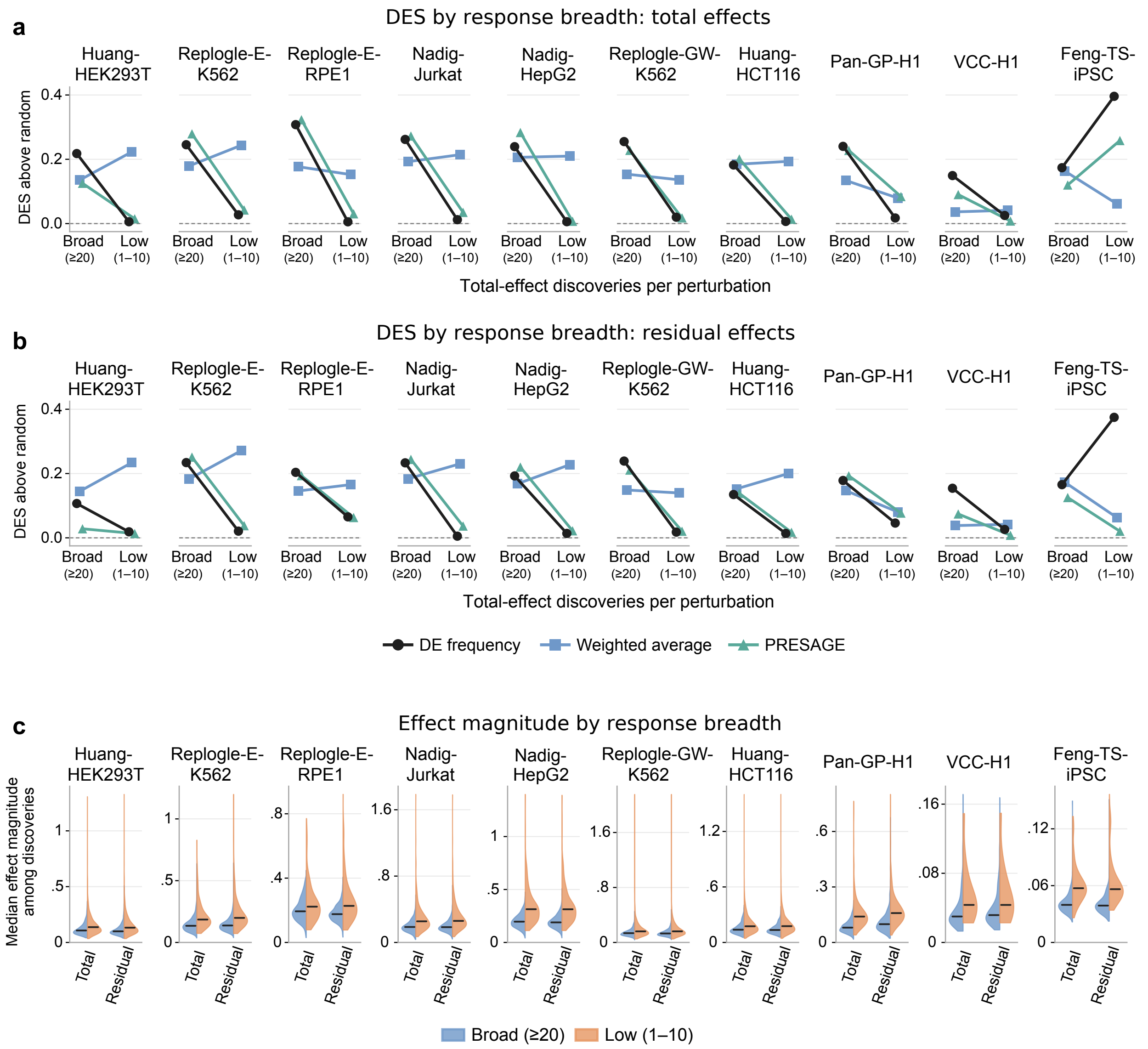


Figure S4: **Discovery prediction and detected-effect magnitudes by response breadth. a,** DES of DE frequency, weighted average and PRESAGE above the random-ranking baseline for total-effect discoveries in broad ($\geq$ 20 total-effect discoveries) and low-discovery (1–10) held-out perturbations in 10 datasets. Points, means over perturbations with $\geq$ 1 residual-effect discovery; the same perturbations are used in b and c. **b,** As in a, for residual-effect discoveries; perturbations are grouped by total-effect discoveries. **c,** Per-perturbation median absolute total effect over total-effect discoveries (Total) and median absolute residual effect over residual-effect discoveries (Residual). Half-violins, broad (blue) and low-discovery (orange) perturbations; black bars, group medians.

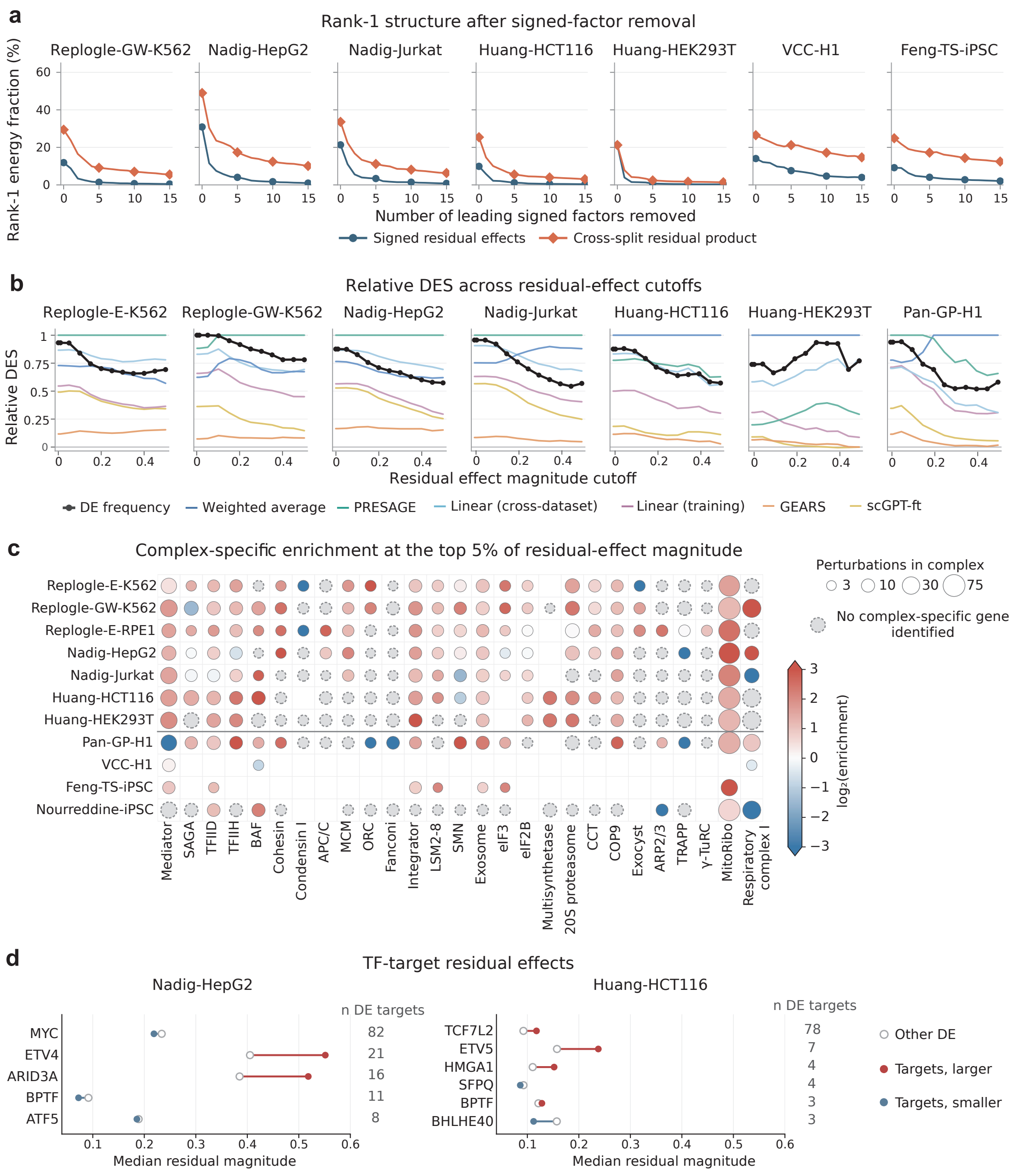


Figure S5: **Magnitude structure and functional organization of residual effects across datasets. a,** Rank-1 energy of signed residual effects and cross-split residual products after removing 0-15 leading signed factors in seven additional datasets not shown in Figure 2c. **b,** Relative DES above increasing residual-effect magnitude cutoffs in seven additional datasets not shown in Figure 2d. **c,** CORUM complex-specific enrichment among the top 5% of discoveries ranked by residual-effect magnitude. Colors show $\log_2$ enrichment capped at $\pm3$; circle size indicates the number of measured perturbation targets in each complex. Gray dashed circles indicate no complex-specific response genes; blank cells indicate $< 3$ measured targets. **d,** Median absolute residual effects for DE targets of HepG2- and HCT116-activated SCENIC+ eRegulons versus other DE genes among transcription-factor perturbations with $\geq 3$ DE targets in Nadig-HepG2 and Huang-HCT116, respectively. Red/blue indicate larger/smaller target medians; numbers indicate DE target counts.

### Supplementary Methods

#### Perturbation-effect estimation

**Regression interpretation.** For the estimator $E_d(p,g)$ defined in the main Methods, let $Y_{is}$ denote the log-normalized expression of cell $i$ in stratum $s$ and let $Z_{is}$ indicate whether the cell carries perturbation $p$. Consider the cell-level regression

$$Y_{is} = \alpha_s + \tau Z_{is} + \varepsilon_{is},$$

where $\alpha_s$ is a stratum-specific intercept. The ordinary least-squares coefficient $\widehat{\tau}$ is exactly $E_d(p,g)$ defined in the main Methods. Thus, the weighting used there is the weighting induced by a regression with stratum-specific intercepts and a single perturbation coefficient across strata.

**Standard errors.** We write $\mathrm{SE}_d(p,g)$ for the standard error of $E_d(p,g)$. Using the cell counts $n_s^{\mathrm{pert}}$, $n_s^{\mathrm{ctrl}}$ and weights $w_s$ defined in the main Methods, let $(s_s^{\mathrm{pert}})^2$ and $(s_s^{\mathrm{ctrl}})^2$ denote the within-stratum sample variances of gene $g$ expression among perturbed and control cells, for strata with at least 2 perturbed or controlled cells, respectively. Assuming that cells are independent within strata and that, for each gene, perturbed and control cells each have a common variance across strata, we estimated the condition-specific variances as

$$\widehat{\sigma}^2_{\mathrm{pert}} = \frac{\sum_{s:\, n_s^{\mathrm{pert}} \geq 2} w_s (s_s^{\mathrm{pert}})^2}{\sum_{s:\, n_s^{\mathrm{pert}} \geq 2} w_s}, \qquad \widehat{\sigma}^2_{\mathrm{ctrl}} = \frac{\sum_{s:\, n_s^{\mathrm{ctrl}} \geq 2} w_s (s_s^{\mathrm{ctrl}})^2}{\sum_{s:\, n_s^{\mathrm{ctrl}} \geq 2} w_s}.$$

The standard error was then estimated as

$$\mathrm{SE}_d(p,g)^2 = \frac{\widehat{\sigma}^2_{\mathrm{pert}} \sum_s w_s^2 / n_s^{\mathrm{pert}} + \widehat{\sigma}^2_{\mathrm{ctrl}} \sum_s w_s^2 / n_s^{\mathrm{ctrl}}}{\left(\sum_s w_s\right)^2}.$$

Strata containing only one cell in a condition could still contribute to the effect estimate but not to estimation of that condition's variance. The pooled variances, and hence the standard error, remained estimable provided that at least one contributing stratum contained at least two perturbed cells and at least one contained at least two control cells. Perturbations for which the required variances could not be estimated were excluded from analyses using standard errors.

#### Stratified Wilcoxon tests

For each perturbation–gene pair, the same strata as in perturbation-effect estimation were used. Within each stratum $s$ containing both perturbed and control cells, expression values were ranked jointly, with average ranks assigned to ties. Let $N_s = n_s^{\mathrm{pert}} + n_s^{\mathrm{ctrl}}$ denote the total number of cells, and $R_s$ the sum of the ranks of the perturbed cells in stratum $s$. Under the null hypothesis,

$$\mathbb{E}_0(R_s) = \frac{n_s^{\mathrm{pert}}(N_s + 1)}{2}.$$

To account for tied expression values, define

$$c_s = 1 - \frac{\sum_j (t_{sj}^3 - t_{sj})}{N_s^3 - N_s},$$

where $t_{sj}$ is the size of the $j$th tied group within stratum $s$. The corresponding null variance is

$$V_s = c_s \frac{n_s^{\text{pert}} n_s^{\text{ctrl}} (N_s + 1)}{12}.$$

The stratum-specific centered rank sums were combined as

$$Z = \frac{\sum_s \{R_s - \mathbb{E}_0(R_s)\}}{\sqrt{\sum_s V_s}},$$

where the sums are over strata containing both perturbed and control cells. The two-sided $P$ value was

$$P = 2\Phi(-|Z|).$$

Equivalently, this procedure sums the stratum-specific Mann–Whitney $U$ statistics and standardizes the sum by its combined null variance. No continuity correction was applied.

#### Noise correction for cross-dataset cosine similarity

For a perturbation $p$ and outcome gene $g$ in dataset $d$, write

$$E_d(p, g) = \theta_d(p, g) + \varepsilon_d(p, g),$$

where $\theta_d(p, g)$ is the underlying perturbation effect and the estimation error satisfies

$$\mathbb{E}\{\varepsilon_d(p, g)\} = 0, \qquad \text{Var}\{\varepsilon_d(p, g)\} = \text{SE}_d(p, g)^2.$$

Assuming independent estimation errors across datasets, the observed squared norm contains an additive noise contribution,

$$\mathbb{E}\left[\sum_g E_d(p, g)^2\right] = \sum_g \theta_d(p, g)^2 + \sum_g \text{SE}_d(p, g)^2,$$

whereas the cross-dataset inner product does not:

$$\mathbb{E}\left[\sum_g E_d(p, g) E_{d'}(p, g)\right] = \sum_g \theta_d(p, g) \theta_{d'}(p, g).$$

Thus, subtracting the summed estimated error variances from each observed squared norm removes its expected additive noise contribution, while no analogous subtraction is required for the cross-dataset inner product. This motivates the corrected squared norm $Q_d(p)$ and cosine similarity defined in the main Methods.

#### Noise correction for signal energy

For the strong-perturbation matrix in dataset $d$, write

$$E_d = \Theta_d + \varepsilon_d,$$

where $\Theta_d$ is the matrix of underlying perturbation effects. For each perturbation $p$, mean-zero estimation error gives

$$\mathbb{E}\Big\{(E_d E_d^\mathsf{T})_{pp}\Big\} = (\Theta_d \Theta_d^\mathsf{T})_{pp} + \sum_g \mathrm{SE}_d(p,g)^2.$$

Thus, subtracting the summed estimated error variances from the diagonal removes the expected noise inflation of each perturbation's squared norm. In particular, for the corrected matrix $\widehat{K}_d$ defined in the main Methods,

$$\mathbb{E}\Big\{\mathrm{tr}(\widehat{K}_d)\Big\} = \|\Theta_d\|_\mathrm{F}^2,$$

so that

$$\mathrm{tr}(\widehat{K}_d) = \|E_d\|_\mathrm{F}^2 - \sum_{p,g} \mathrm{SE}_d(p,g)^2$$

estimates the total squared signal energy.

For two distinct perturbations $p \neq p'$, the corresponding off-diagonal entry satisfies

$$\mathbb{E}\Big\{(E_d E_d^\mathsf{T})_{pp'}\Big\} = (\Theta_d \Theta_d^\mathsf{T})_{pp'} + \sum_g \mathrm{Cov}\{\varepsilon_d(p,g), \varepsilon_d(p',g)\}\,.$$

Therefore, if estimation errors are uncorrelated across perturbations, the mean of $\widehat{K}_d$ is $\Theta_d\Theta_d^\mathsf{T}$. We used $\lambda_{\max}(\widehat{K}_d)/\,\mathrm{tr}(\widehat{K}_d)$ as a noise-corrected plug-in summary of the fraction of signal associated with the leading rank-one structure.

#### Feature-gene selection

For the feature-gene sensitivity analysis in Supplementary Fig. S1a, each outcome gene $g$ was scored by the variance of its signed total effects across all QC-passing perturbations,

$$V_{dg} = \mathrm{Var}_{p\in\mathcal{P}_d:\,p\neq g}\{E_d(p,g)\},$$

where $\mathcal{P}_d$ is the set of all perturbations retained after preprocessing. The direct-target entry was treated as missing rather than zero, and missing effect estimates were ignored. The implementation used population variance. Genes were ranked within each dataset by $V_{dg}$, and the 2,000 highest-ranking outcome genes were retained as feature genes. Feature scoring used total effects from all perturbations and was not restricted to strong perturbations or statistically significant effects. The selected gene set was held fixed after removal of the leading singular component.

#### Sensitivity analyses for cross-dataset similarity

Supplementary Fig. S1a examined sensitivity to the perturbation and outcome-gene sets used for cross-dataset similarity. The primary analysis used strong perturbations and all outcome genes retained after preprocessing. In the left panels, total- and residual-effect similarities were recomputed using all perturbations and the same outcome-gene set. In the right panels, similarities were recomputed using strong perturbations and the 2,000 feature genes defined above. In all cases, the leading singular component was the component fitted once to the full perturbation-by-outcome-gene total-effect matrix for each dataset and was not refitted after subsetting. Strong-perturbation and feature-gene selections were defined from total effects and held fixed for the corresponding residual analyses. All other pairwise overlap, direct-target exclusion, noise-correction, and filtering steps followed the primary cross-dataset similarity analysis.

To assess sensitivity to the noise correction (Supplementary Fig. S1b), we compared the similarities of the original VCC-H1 dataset and a version downsampled in both UMI depth and cell number with each other dataset. Starting from raw counts, we randomly kept 16 of the 48 batches and then randomly sampled up to 100 cells per perturbation, keeping all control cells in these batches. The counts for each cell were then randomly subsampled by drawing 13,000 UMIs from each cell without replacement (cells with fewer were unchanged), which is close to the median library size of Replogle-E-K562. These steps matched the cell numbers per perturbation and the library sizes of the essential-gene screens, while keeping control cells from the same batches for each perturbation. The downsampled dataset was then processed as the other datasets.

Supplementary Fig. S1c assessed concordance of perturbation strength across datasets. For each dataset pair, we computed the Spearman correlation between the L1 perturbation-strength scores $S_{dp}^{\mathrm{L1}}$ across perturbations shared by the two datasets.

#### Leading-direction concordance

Supplementary Fig. S1d assessed concordance of the leading response direction across datasets. For dataset $d$, let $u_{1d}$ and $v_{1d}$ denote the leading left and right singular vectors, respectively, from the SVD fitted to all perturbations and all primary outcome genes. For a pair of datasets $d$ and $d'$, let $P$ and $G$ denote their shared perturbations and outcome genes, and let $u_{1d,P}$ and $v_{1d,G}$ denote the corresponding restricted vectors. We computed

$$C_{dd'} = \frac{u_{1d,P}^{\top} u_{1d',P}}{\|u_{1d,P}\|_2 \, \|u_{1d',P}\|_2} \cdot \frac{v_{1d,G}^{\top} v_{1d',G}}{\|v_{1d,G}\|_2 \, \|v_{1d',G}\|_2}.$$

Thus, concordance is the product of the cosine similarities between the perturbation-side and gene-side leading singular vectors after restriction to their shared entries. The SVD was not refitted for individual dataset pairs.

#### Prediction analyses

**Implementation of comparison methods.** GEARS and scGPT were trained separately on each target dataset, with scGPT fine-tuned on the target-dataset training perturbations. For Linear (training), the perturbation representation was constructed from the target-dataset training effects following the same linearModel procedure as in the original paper[8]. For Linear (cross-dataset), perturbation effects were first averaged across the same reference datasets from other cell lines available to weighted average, and the resulting effects were used to construct the SVD-based perturbation representation following the original paper[8]. For PRESAGE, we used PRESAGE (+Perturb-seq), augmenting its knowledge-source embeddings with Perturb-seq embeddings derived from the same reference datasets available to weighted average, following the same procedure as the original paper[9]. To reduce the heavy computation for GEARS and scGPT in Genome-wide screens, control cells were randomly subsampled, capped at 50,000 in Feng-GW-iPSC, Huang-HCT116, Huang-HEK293T and Nourreddine-iPSC, and training perturbations were further randomly subsampled to one third in Huang-HCT116, Huang-HEK293T and Nourreddine-iPSC.

**Weighted mean squared error.** For perturbation $p$, let $z_{pg}$ denote the stratified Wilcoxon $z$-statistic for outcome gene $g$. Following the weighted-MSE construction proposed in the original paper[10], we defined

$$a_{pg} = \left( \frac{|z_{pg}| - \min_{g'} |z_{pg'}|}{\max_{g'} |z_{pg'}| - \min_{g'} |z_{pg'}|} \right)^2, \qquad w_{pg} = \frac{a_{pg}}{\sum_{g'} a_{pg'}},$$

and

$$\mathrm{wMSE}(p) = \sum_g w_{pg} \left\{ E(p,g) - \widehat{E}(p,g) \right\}^2.$$

We reported the relative reduction in wMSE versus the mean baseline,

$$\frac{\mathrm{wMSE}_{\mathrm{mean}} - \mathrm{wMSE}_{\mathrm{method}}}{\mathrm{wMSE}_{\mathrm{mean}}},$$

using method and mean-baseline errors evaluated on the same test fold.

**Perturbation discrimination score.** For a test fold containing $n$ perturbations, we ranked the observed effects of all test perturbations by increasing cosine distance from the prediction for each perturbation $p$. If $r_p$ is the zero-based rank of the observed effect corresponding to $p$, then

$$\mathrm{PDS}(p) = 1 - \frac{r_p}{n-1}.$$

Thus, $\mathrm{PDS}(p) = 1$ when the correct observed perturbation is closest to its prediction. The mean baseline gives the same predicted vector for every test perturbation; when distances are distinct, the correct perturbations occupy ranks $0, \dots, n-1$ once each and its fold-average PDS is therefore 0.5. We report PDS $-$ 0.5, so that the mean baseline corresponds to zero on this scale.

**Residual cosine similarity.** Within each test fold, we removed the leading rank-one component separately from the observed and predicted perturbation-effect matrices. If

$$E = U \Sigma V^{\mathsf{T}}, \qquad \widehat{E} = \widehat{U} \widehat{\Sigma} \widehat{V}^{\mathsf{T}},$$

we defined

$$R = E - \sigma_1 u_1 v_1^{\mathsf{T}}, \qquad \widehat{R} = \widehat{E} - \widehat{\sigma}_1 \widehat{u}_1 \widehat{v}_1^{\mathsf{T}}.$$

For each perturbation, residual cosine similarity was calculated between the corresponding rows of $R$ and $\widehat{R}$ over all evaluated outcome genes and, separately, over the top 20 and top 100 differentially expressed genes in the target dataset, ranked by stratified Wilcoxon $P$ value. The mean baseline has identical prediction vectors across perturbations, so its prediction matrix is rank one and its residual prediction is zero.

**Metric aggregation.** Perturbation-level metrics were averaged over test perturbations within each fold and summarized as mean ± s.d. across the five folds.

#### Total-effect conjunction

Supplementary Fig. S3a compares two conjunction. The residual-effect conjunction defined residual discoveries (Methods). The total-effect conjunction applies the same criterion to the total effect,

$$p_{pg}^{\mathrm{conj},E} = \max\left\{p_{pg}^{\mathrm{W}},\ 2\Phi\left(-\frac{|E(p,g)|}{\mathrm{SE}(p,g)}\right)\right\},$$

with BH adjustment within each perturbation and a threshold of 0.05. For each criterion, we report the percentage of total-effect discoveries it retains.

#### Gene-responsiveness hierarchy and discovery prediction

For visualization of the held-out discovery matrices, perturbations were ordered by their discovery fraction and genes by the corresponding training-derived discovery-frequency ranking. In each split, the ordered perturbations and genes were each divided into 20 quantile groups, producing a $20 \times 20$ grid. Each grid entry was the mean discovery indicator among perturbation–gene pairs in the corresponding pair of quantile groups. These $20 \times 20$ matrices were then averaged across repeated splits, providing a common representation despite differences in the identities and ordering of held-out perturbations across splits.

Capture curves were evaluated only for held-out perturbations with at least one discovery under both the total- and residual-effect definitions, so that the two curves were based on the same held-out perturbations.

#### Progressive factor removal and cross-split residual-product analysis

The cross-split construction was used to distinguish structure in underlying residual-effect magnitude from apparent structure induced by estimation noise. To motivate this construction, suppose that an estimated residual effect after removal of the first $k$ signed factors can be written as

$$R_k(p,g) = \theta_k(p,g) + \varepsilon_k(p,g),$$

where $\theta_k(p,g)$ denotes the underlying residual effect and $\varepsilon_k(p,g)$ is a mean-zero estimation error with variance $\sigma_k^2(p,g)$. Directly analyzing squared observed effects does not isolate the magnitude of the underlying signal, since

$$\mathbb{E}\Big[R_k(p,g)^2\Big] = \theta_k(p,g)^2 + \sigma_k^2(p,g).$$

Thus, structured heterogeneity in estimation variance across perturbations and outcome genes can itself generate structure in the matrix of observed squared effects.

Absolute observed effects have an analogous problem. For example, consider the null case $\theta_k(p,g) = 0$ and Gaussian estimation errors $\varepsilon_k(p,g) \sim N\{0, \sigma_k^2(p,g)\}$. Then

$$\mathbb{E}|R_k(p,g)| = \sqrt{\frac{2}{\pi}}\,\sigma_k(p,g).$$

Consequently, even when the true residual-effect matrix is identically zero, systematic variation in estimation precision can produce low-rank structure in observed effect magnitudes. In particular, if the noise standard deviations factorize as $\sigma_k(p,g) = a_p b_g$, then

$$\mathbb{E}|R_k(p,g)| = \sqrt{\frac{2}{\pi}}\, a_p b_g,$$

which is exactly rank one despite the absence of any true residual effects. The same example gives

$$\mathbb{E}\Big[R_k(p,g)^2\Big] = a_p^2 b_g^2,$$

which is also rank one.

We therefore estimated residual effects independently using two disjoint halves of the cells. For each random split, cells carrying each perturbation and control cells were divided into two halves, effects were estimated separately, and the leading signed components were fitted and removed separately within each half. Write the two residual estimates as

$$R_k^{(s)}(p,g) = \theta_k(p,g) + \varepsilon_k^{(s)}(p,g), \qquad s = 1,2.$$

We then formed the cross-split product

$$M_k(p,g) = R_k^{(1)}(p,g) R_k^{(2)}(p,g).$$

Under mean-zero estimation errors that are independent across the two cell halves,

$$\begin{aligned}\mathbb{E}[M_k(p,g)] &= \mathbb{E}\Big[\{\theta_k(p,g) + \varepsilon_k^{(1)}(p,g)\}\{\theta_k(p,g) + \varepsilon_k^{(2)}(p,g)\}\Big] \\ &= \theta_k(p,g)^2.\end{aligned}$$

Thus, the cross-split product targets squared underlying residual-effect magnitude without the additive estimation-variance contribution present in squared effects estimated from a single sample. In particular, when $\theta_k(p,g) = 0$, its expectation is zero regardless of the entry-specific estimation variance. Individual cross-products remain noisy and can be negative, but averaging across independent splits reduces this variability without introducing the systematic positive contribution created by taking absolute values or squares of noisy estimates.

#### Extreme-entry capping for rank-one energy

For each dataset and factor-removal depth $k$, the half-specific residual matrices were first averaged to obtain the signed-residual matrix, and the corresponding cross-split product matrices were averaged to obtain the residual-magnitude matrix. Extreme-entry capping was applied only to these final averaged matrices, before calculation of the rank-one energy; it did not affect the half-specific SVD fits used for factor removal.

For each averaged matrix $X$, we calculated the 99.99th percentile of the absolute values of its eligible non-target entries,

$$c_X = Q_{0.9999}\left(\{|X_{pg}|\}\right),$$

and replaced each entry by

$$X_{pg}^{\text{cap}} = \text{sign}(X_{pg}) \min\{|X_{pg}|, c_X\}.$$

The cutoff was calculated across the full matrix and separately for each dataset, factor-removal depth, and matrix type. Target-gene entries were excluded when determining the cutoff. Rank-one energy was then calculated from the capped matrix as

$$F_1(X^{\text{cap}}) = \frac{\sigma_1(X^{\text{cap}})^2}{\|X^{\text{cap}}\|_{\text{F}}^2}.$$

This capping was used as a robustness adjustment because a very small number of extreme entries can account for a substantial fraction of the total squared energy and thereby dominate the leading singular component. This issue is especially relevant for the cross-split product matrices, whose expected signal is the squared underlying effect, so genuinely large effects can produce disproportionately large entries. The procedure preserves entry signs and alters only the extreme absolute-magnitude tail. It is not a noise correction and may also attenuate genuinely large effects; accordingly, the reported rank-one energy fractions describe the capped matrices.

#### Response-breadth analysis

Response breadth was summarized by the number of discoveries of total effects for each perturbation. We compared prediction performance and detected-effect magnitudes between perturbations with broad responses and those with few discoveries. Broad and low-discovery perturbations were defined as having at least 20 and 1-10 total-effect discoveries, respectively, and the same perturbation groups were used for analyses of residual discoveries.

Detected-effect magnitude was summarized by the median $|E(p,g)|$ over total-effect discoveries and, separately, the median $|R(p,g)|$ over residual discoveries. Prediction performance was evaluated using DES for the corresponding total- and residual-effect discovery sets (Fig. S4).

#### Protein-complex enrichment

We used 27 human protein complexes from CORUM 5.0: Mediator, SAGA, TFIID, TFIIH, BAF, cohesin, condensin I, APC/C, MCM, ORC, the Fanconi anaemia core complex, Integrator, LSm2–8, SMN, the exosome, eIF3, eIF2B, the multisynthetase complex, the 20S proteasome, CCT, the COP9 signalosome, the exocyst, Arp2/3, TRAPP, $\gamma$-TuRC, the 55S mitochondrial ribosome, and respiratory complex I. Within each dataset, we considered residual discoveries from perturbations targeting members of these complexes and retained complexes with at least three measured perturbation targets.

For each eligible complex $A$ and outcome gene $g \notin A$, we quantified whether $g$ was preferentially discovered following perturbations of members of $A$. Let $Y_{pg} = \mathbf{1}\{g \in D'_p\}$ denote the residual-discovery indicator. We first defined

$$s_{Ag} = \frac{1}{|P_A|} \sum_{p \in P_A} Y_{pg} - \frac{1}{|P_{\text{C}} \setminus P_A|} \sum_{p \in P_{\text{C}} \setminus P_A} Y_{pg},$$

where $P_A$ is the set of measured perturbations targeting members of complex $A$ and $P_{\text{C}}$ is the set of all perturbations targeting members of the considered complexes.

Because some perturbations produce many discoveries and some outcome genes are broadly responsive across perturbations, we additionally removed the discovery frequency expected from these two marginal tendencies. Among complex-targeting perturbations, let $d_p$ denote the fraction of outcome genes discovered for perturbation $p$, $c_g$ the fraction of perturbations for which gene $g$ was discovered, and $\rho$ the overall discovery fraction. We defined the adjusted discovery indicator

$$Y'_{pg} = Y_{pg} - \frac{d_p c_g}{\rho},$$

and calculated the corresponding within-complex enrichment

$$s'_{Ag} = \frac{1}{|P_A|} \sum_{p \in P_A} Y'_{pg} - \frac{1}{|P_\mathrm{C} \setminus P_A|} \sum_{p \in P_\mathrm{C} \setminus P_A} Y'_{pg}.$$

Outcome gene $g$ was defined as complex-specific for $A$ if $s_{Ag} > 0.25$, $s'_{Ag} > 0.25$, the two-proportion $z$-statistic corresponding to $s_{Ag}$ exceeded 2.5, and $g$ was discovered for at least three perturbations targeting members of $A$.

To evaluate whether these complex-specific responses were enriched among larger residual effects, we pooled residual discoveries from perturbations targeting eligible complexes. Perturbation-gene pairs for which the perturbed gene and outcome gene belonged to the same complex were excluded. For each eligible complex $A$, let $\mathcal{G}_A$ denote its set of complex-specific outcome genes. A residual discovery $(p, g)$ was labeled complex-specific if there existed an eligible complex $A$ such that the target of perturbation $p$ belonged to $A$ and $g \in \mathcal{G}_A$.

We ranked the remaining residual discoveries by $|R(p, g)|$. For the top $q$% of discoveries in this ranking, enrichment was defined as the fraction labeled complex-specific divided by the corresponding fraction among all residual discoveries. Enrichment was evaluated for $q$ from 100 to 1 and summarized across datasets by the median. Complex-level enrichment was additionally summarized for each complex separately at $q = 5\%$ (Fig. S5c).
